# Structure of bacteriophage P1 head provides insight into capsid polymorphism

**DOI:** 10.1101/2025.11.26.689930

**Authors:** Anindito Sen, Tsukasa Nakamura, Genki Tarashi, Veer Bhatt, Kyungho Kim, Shankararaman Chellam, Le Tran, Daisuke Kihara

## Abstract

The highly regulated assembly of different proteins forming mature P1 myophage capsid show a unique polymorphism where the virion population is primarily divided in brackets of two different capsid sizes. The contrasting capsids, sized 95nm (capsid_L), triangulation number (T)=13 and 68nm (capsid_S) with T=7, both in Dextro format and have the same protein forming the phage head with similar intra and inter capsomeric interactions. Comparative study of the electron density maps of the capsids reveals the presence of protein appendages DarA and Hdf below the 5-fold symmetric region inside capsid_L that anchor the phage dsDNA to the capsid-shell. Such densities are also noticed in large capsids depleted off their dsDNA but not observed in capsid_S. Deficiency of these appendages in capsid_S results in a smaller sized dsDNA packed inside capsid_S. Despite missing out on nearly 60% of the phage dsDNA virions with capsid_S has all the essential genes responsible for the formation of a fully mature thermodynamically stable virion particle with structurally identical tail and baseplate as of capsid_L. Co-existence of both stable conformation in a single lysate is a unique phenomenon in the phage community and suggest that DarA and Hdf play an important role in phage capsid morphogenesis.

---

In a normal P1 phage lysate, nearly 80% of the virion particles have capsid_L while the rest 20% are capsid_S with both the capsids possessing tightly packed dsDNA inside them (Fig. 1A) (1). While a third type of P1 virion with capsid size even smaller than capsid_S was reported (∼45 nm) but not visualized in our specimen (SI) (2,3). The atomic model of the main capsid protein (MCP) gp23 of P1, is determined employing icosahedral reconstruction of phage-head images from cryo-TEM data followed by computation image analysis (SI). The MCP of P1 phage has a polypeptide fold similar to gp23 capsid protein of bacteriophage T4 (Fig. 1B) (4,5). It has three domains termed axial (A), peripheral (P) and insertion (I). There are three loops AP, P, P-loop short helix (Psh) and a backbone helix. Each of these elements play a significant role in the inter (boxed in blue) and intra (boxed in red) capsomer interactions, described later (5). The variation in the motif of P1 capsid size can be explained by elaborating the difference among the asymmetric units of capsid_L and capsid_S, where the latter is missing 6 molecules of gp23, resulting a total of 360 capsomers less than capsid_L (Fig. 1A, C). The core volume capsid_L is 107 x 10^3^ nm^3^ that encapsulates a 32,500 nm long (93-95 kbp) dsDNA resulting in a 0.3 bp/nm^3^ packing factor indicating that the P1 genome (dsDNA) is in a reasonably condensed state inside the capsid (6). The spacing between the icosahedral symmetrized dsDNA in the electron density maps for either L or S capsids are found to be roughly 2.7 nm suggesting they have a same packing factor (Fig. 1F(i)). The dsDNA length enclosed inside capsid_S of volume of approximately 40.0 x 10^3^nm^3^ is computed approximately to a length of 12,150 nm that corresponds to nearly 37.7 kbp, which is around 40% of the actual phage dsDNA (2). A single protein gp23 is found to frame the shell of both types of the capsids and the size dissimilarity does not alter the nature of molecular interactions among MCP molecules. However, the N-terminal region of the gp23 capsomer of a penton is found to have an arc-like conformation when compared to a hexon counterpart, indicating the adaptiveness of the molecule to alter its conformation for fabricating stable intra and inter capsomer interactions (Fig. 1D). In a hexon, intra molecular interactions take place among the A-A domains of two adjacent capsomers through strong Hydrogen (H) bond among residues Arg 405 to Asp 425 and at the center, strong electrostatic bonds are found among the A-loops involving residues Lys551 and Asp563 (Fig. 1E (i,v)). Further interactions take place among the P-P-I domains of the two adjacent capsomers, where residues Ile310 with Ala327 and Gln323 with Glu297 form ‘H’ bonds (Fig. 1E(ii)). Intermolecular interactions among two neighboring hexon molecules occur at the 2-fold axis where interaction Psh loop and P domain of neighboring subunits takes place (Fig. 1E(iii)). For a hexon-penton intermolecular interlinkage, the Psh loop and the tip of the P-domain of the hexon capsomer form bonds with the P domain of the one of penton capsomer and I domain linker of the adjacent penton capsomer, respectively suggesting a stronger hold between the symmetry mismatched 5- and 6-fold regions that helps structurally stabilizing the bend regions of the capsid (Fig. 1E(iv)). The penton intra-capsomeric interactions are different than its hexon counterpart. Starting at the 5-fold tip regions, the 5 A-loops of the are somewhat spaced-out due the absence of a capsomer (when compared the same region of the hexon) causing electrostatic interactions to occur between Lys 550 and Asp560 that are positioned below the tip region (Fig. 1E(vi)). The P-P-I domain contacts, seen in the intra-penton capsomer interaction is common in the intra-hexon counterpart (Fig. 1E(viii)). In both penton and hexon a set of ‘H’ bonds among the **β**-strands of the P domain providing structural stability to the capsid (Fig. 1E(vii)).

**Figure 1.**
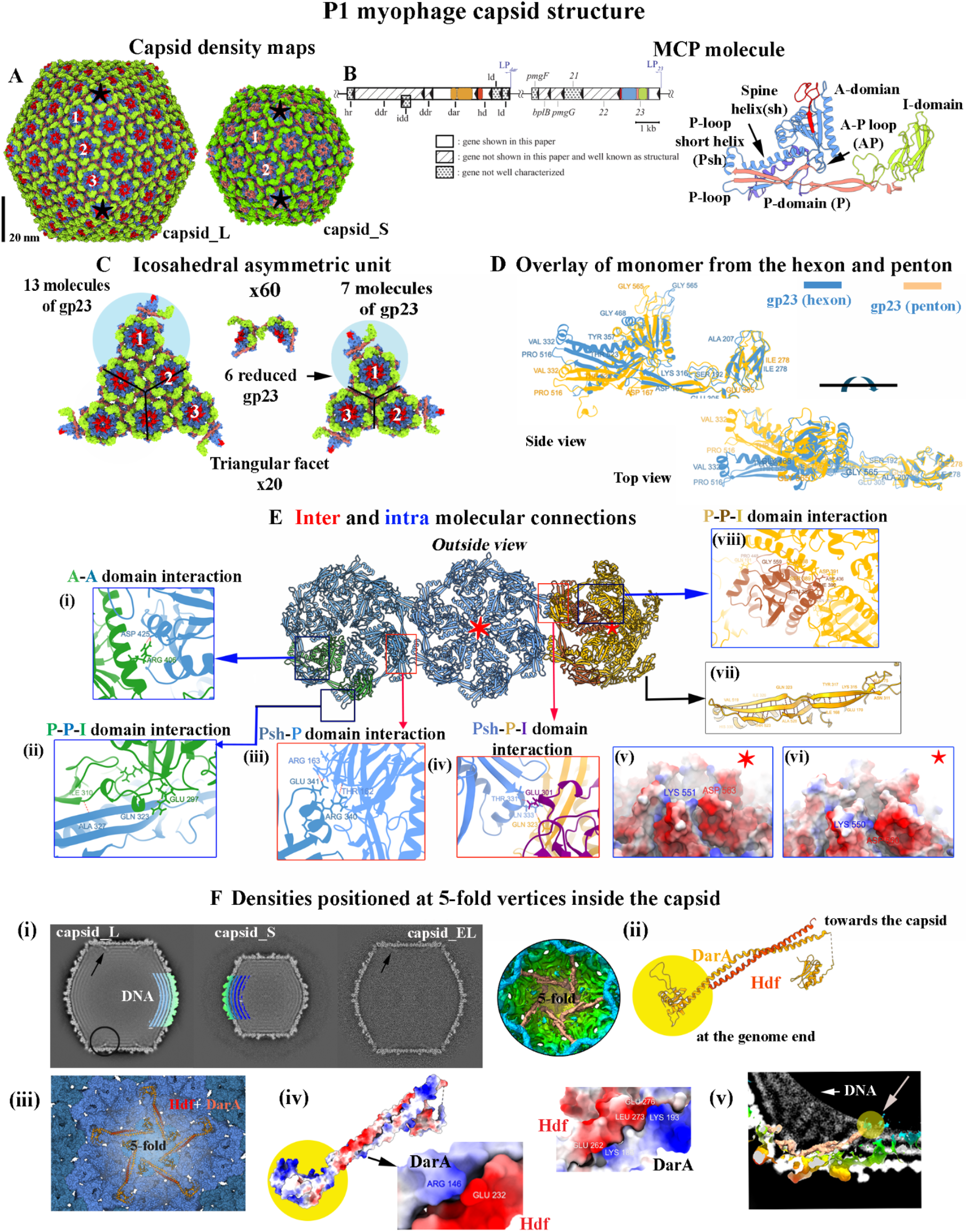
P1 myophage capsid. (A) electron density maps of capsid_L and capsid_S at resolution 3.8Å and 4.17Å at Fourier Shell Correlation (fsc) cutoff of 0.143, respectively. The black stars represent the pentons of the capsid surface. The numbers 1, 2 and 3 represent hexons, a count for computing the T number of the capsids, 13 for capsid_L and 7 for capsid_S. (B) Gene Diagram and atomic structure of P1 MCP. Boxes with internal triangles represent the positions on the P1capsid protein genome and orientations of the genes with their names. Colors of the boxes correspond to the colors of the parts of MCP ribbon structure, shown next to the gene diagram. Gene boxes without hatch are genes shown in this paper. Boxes with slashes are genes not shown in this paper and are well known as structural proteins in previous studies (2). Boxes with dots are genes that are not well characterized. Atomic structure of MCP of P1 capsid. The A domain along with the spine helix (blue), AP loop (blue), P domain and I-domain linker (salmon), I-domain linker, I domain (green) are shown. The overall coloration provides a clear view of the distribution of the pentons and hexons on the capsid surface (A) and the icosahedral asymmetric unit in (C). A triangle facet (20 overall in the entire icosahedral) of both the capsids is shown with the asymmetric units highlighted by blue background. Capsid_L has 13 monomers. Eliminating 6 monomers, (shown in the center), results in the asymmetric unit of capsid_S that has T=7. (D) Top and side views of the overlay of the gp23 monomer of a penton and hexon along with some representative residues showing the A, P domains and the associated loops of the penton have adapted to an arc-shape. (E) Ribbon diagram of 2 hexons and a penton and their corresponding intra (blue boxed) (i, ii & viii) and inter (red boxed) (iii,iv) capsomeric interactions. A single capsomer is colored in green in one of the hexon and another as brown in the penton to illustrate the interactions. The electrostatic interactions among the capsomers at the tip of the hexon and pentons are shown in (v) and (vi) respectively. The bonds between the parallel ? strands providing structural rigidity of the capsid. (F) Densities at the 5-fold regions inside the capsid. (i) central slices of capsid_L, capsid_S and capsid_EL. The protein appendages at one of the 5-fold region are circled. A portion of the reconstructed electron density maps with the dsDNA shells overlaid on the slices. (ii) Iso-surface representation of the 5-fold region with the densities representing DarA and Hdf colored in brown. A large segment of density representing dsDNA is computationally removed for better visualization. Atomic structure of the densities: the large helix (colored gold) with globular domains is the DarA protein while the smaller helix (colored red) is Hdf protein. The yellow-colored background globular domain in (ii) is primarily electropositive in nature shown in (iv) and interacts with the phage DNA, pointed by an arrow in (v), which is an overlay of a cropped slice of density map of capsid_L at the 5-fold region with the dsDNA computationally removed on the central slice of the same density map.. An overview of the 5-fold region with the atomic structures of the capsid and DarA with Hdf is shown (iii). Both DarA and Hdf are electrostatically held to each other with opposite charged residues engaging with each other at different points shown in (iv).

A comparative study of the inside of capsid density maps revealed presence of thin protein appendages at the 5-fold regions that anchor the tightly packed dsDNA to the capsid shell in the capsid_L but is absent in capsid_S. However very weak densities, resembling these appendages were observed in the density map of empty capsids (capsid_EL). The plausible reason of these densities appearing feeble is possibly due to the loss of the dsDNA that releases one end of the appendages free, their density in the reconstructed map is averaged over a region and appears feeble (Fig. 1F(i)). Additionally, the loss of the dsDNA perhaps has resulted in loss of some of the appendages dissociating from the capsid shell. Modelling of these appendages in the computed electron density maps of capsid_L revealed that these two appendages are long ‘**α**’ helices that are products antirestriction genes *darA* and *hdf* (Fig. 1E(ii-iii)) (SI). The longer helix is DarA protein with 2 globular densities on either end and the smaller helix is Hdf protein, both are held together by strong electrostatic forces with the N-terminal region of DarA is the globular domain predominantly positively charged, interacts with the negatively charged dsDNA and holds the genome to the inner portion of 5-fold region of the capsid (Fig. 1F(iv-v)). The strong electrostatic charges between the two, aid in this process and the Hdf protein appears to provide a mechanical support to the DarA to act like a molecular anchor bringing mechanical stability to the packed phage dsDNA inside the capsid (2). Despite missing out on nearly 60% of dsDNA of the capsid_L, the dsDNA of capsid_S has all the essential genes responsible for the formation of a fully mature stable virion particle. Our results advices that P1 capsid in-it-self may not be architecturally rigid enough to encapsulate dsDNA of size larger than 37.7 kbp thus forming capsid_S. However, with the mechanistic support offered by the DarA and Hdf, P1 packages nearly 2.5 times of the dsDNA in capsid_S thus forming capsid_L. What additional structural and/or functional characteristics the additional 60% of the dsDNA brings to the P1 virion capsid_L remains to be investigated.

## Acknowledgements

The authors thank Dr. Tom Goddard of University of California San Francisco for his significant help in developing the movie showing the step infection process. The authors also thank Dr. Avery L. McIntosh and Dr. Lawrence Griffin of Texas A&M University for their support in this work. The authors also thank and acknowledge Mr. Gregory G. Servos of OvationData for his assistance with the project.

## Authors’ contribution

Conceptualization: AS. Methodology: AS. Specimen Preparation: AS and KK. Data collection: VB and AS. Computation image processing: AS, LT. Model building: TN, GT. Manuscript preparation: AS, TN, GT, DK, SC.

## Data Availability

Electron density maps for capsid_L: EMD-27716, capsid_S: EMD-74053

## PDB accession numbers

9zdn.

## Supplementary Information (SI)

## Materials and Methods

### Virus Propagation and Purification

Bacteriophage P1 (ATCC 25404-B1) was propagated using a host bacterium Escherichia coli following a confluent lysis method (12). The host was grown in Luria-Bertani (LB) broth (10 g/L tryptone, 5 g/L yeast extract, 10 g/L NaCl, and 10 mM MgSO4) until its optical density at 600 nm wavelength (OD600) reached 0.35 (HACH DR6000), which was then inoculated with P1 phages at multiplicity of infection of 1 in presence of 5 mM CaCl2. A 100 µL of the mixture was added to 10 mL soft LB agar (LB broth amended with 5 g/L agar and 5 mM CaCl_2_) and poured on a LB agar plate (LB broth amended with 15 g/L agar and 5 mM CaCl2). The plate was incubated at 37 °C until a clear phage lawn appeared because of confluent lysis. The lawn was collected and centrifuged (10,000 g for 10 min at 4 °C) after which supernatant was recovered. For further purification, supernatant was filtered with 0.45 µm polyethersulfone (PES), pelletized (15,000 g for 24 hours at 4 °C), and resuspended in 3 mL SM buffer (50 mM Tris-HCl, 100 mM NaCl, 8 mM MgSO4, pH 7.3) overnight at 4 °C. Afterwards, the resultant was passed through CsCl layers (1.3-1.6 g/cm3 prepared in SM buffer, 104,000 g for 24 hours at 4 °C). Phage band was recovered, pelletized (15,000 g for 24 hours at 4 °C), and resuspended in 1 mL SM buffer overnight at 4 °C. The titer of the final stock was found to be approximately 10^11^ PFU/mL based on the double agar layer method.

### Sample preparation for cryo-TEM and data collection

Copper 300-mesh grids supported with a formvar layer and coated with an ultrathin layer of carbon were prepared by glow-discharging for 1 min at 10 mA under vacuum (PELCO 118 easyGlow). About 5μlts of P1 virus stock (10^12^ particles/ml) suspension were directly mounted on the grids followed by negative staining with 2% uranyl acetate solution (pH not adjusted). Grids were screened using ThermoFisher TF20 operating at an accelerating voltage of 200 keV equipped with Gatan K2 direct detector and Gatan Tridiem GIF-CCD camera for quality and concentration of the virion particles. For cryo-TEM, approximately 3.0 µL of purified virus stock was loaded on holey carbon film grids with 2.0 µm holes (C-flatTM 124, Electron Microscopy Sciences), blotted in an environmentally controlled chamber at 80% humidity, and vitrified by sudden plunging the grids into liquid ethane at -180 °C (EM GP2 Automatic Plunge Freezer, Leica). Prepared cryo-samples were imaged with ThermoFisher TF20 operating at an accelerating voltage of 200 keV equipped with Gatan K2 direct detector, Gatan Tridiem GIF-CCD camera and ThermoFisher Glacios cryo-TEM equipped with an autoloader and Falcon 4 direct electron detector (DED). The grids carrying the vitrified sample were imaged on Glacios cryo-transmission electron microscope (ThermoFisher Scientific) equipped with a field emission gun operated at 200 kV. The microscope was aligned at parallel illumination. Grids were screened for ice quality and particle distribution. For the most promising grids, a total of 4200 movies were recorded in EER format using a Falcon 4 DED at a nominal magnification of 73,000X. The movies were acquired within a defocus range of -0.5 μm to -3.0 μm in an automated fashion using EPU software (ThermoFisher Scientific). Each movie was recorded in counting mode using a total exposure dose of 30 e/Å^2^ at a low dose rate of 3.2 e/Å^2^/sec to avoid coincidence loss. Additional details of data acquisition and processing are summarized below.

**Table S1.** Glacios cryo-TEM set up and data collection parameters.

|  |  |
| --- | --- |
| Nominal magnification | 73,000x |
| Apertures ($\mu\text{m}$) | C2 aperture = 50 $\mu\text{m}$; Objective aperture: n |
| Spot size | 5 |
| Pixel size ($\text{\AA}/\text{pixel}$) | 1.3889 |
| Total dose ($\text{e}/\text{\AA}^2$) | 30 |
| Total number of EER frames per movie | 2,267 |
| Exposure time (sec) | 9.4 |
| Defocus | -0.5 to -3.0 $\mu\text{m}$ |

**Figure S1.**
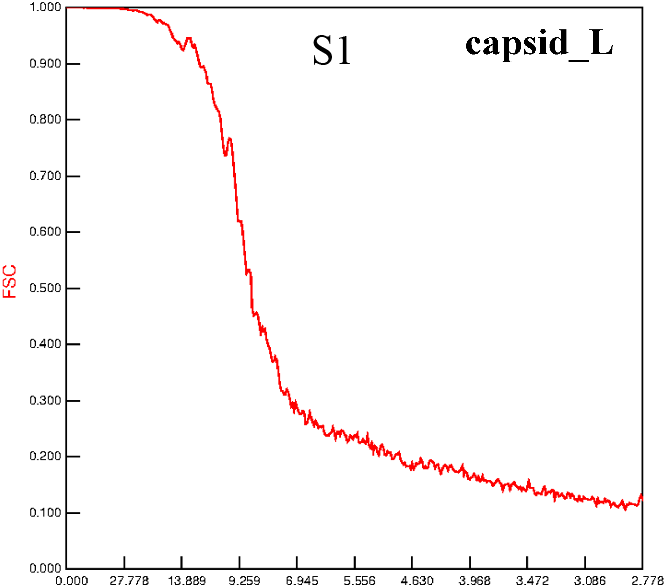

**Figure S2.**
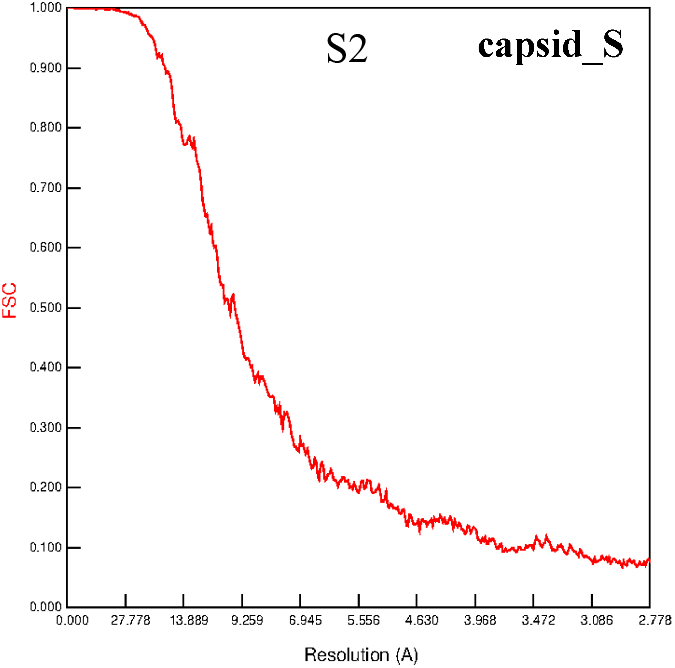

**Figure S3.**
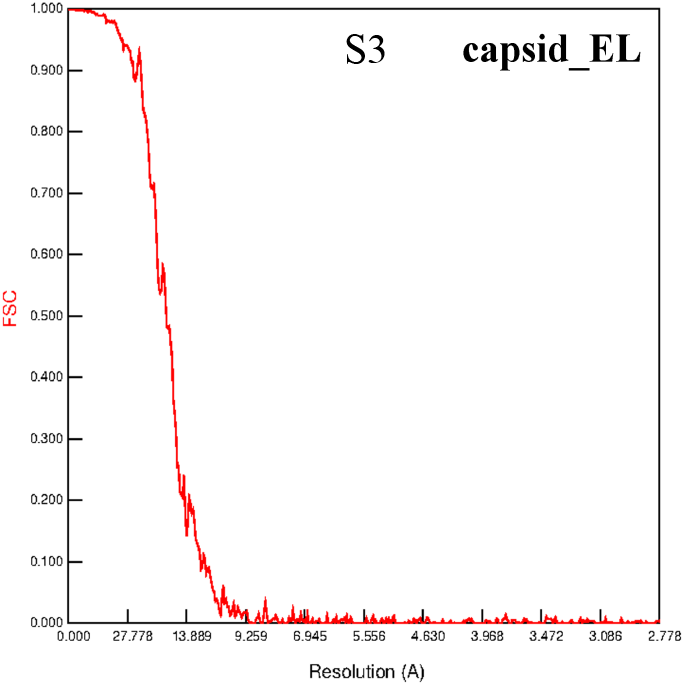

### Computational Image processing

The frames of the movies of the TF20 cryo data recorded were aligned. Movies with frames that showed poor alignment were left out. A total of 7437 capsids were computationally boxed out. The particles were classified primarily in three groups capsid_L, capsid_S and capsid_EL using EMAN2. While capsid_S particles were well separated from the other two types, multiple rounds of classifications were performed to sort out particles belonging to capsid_L and capsid_EL categories. Each of those particle groups were then subjected to single particle computational analysis, with icosahedral symmetry imposed and density maps with resolution 11.2Å, 14.6Å and 21.2Å at a Fourier shell correlation (fsc) cutoff of 0.143 were obtained using BSOFT package (7). Two synthetic icosahedral density maps similar to the sizes of capsid_L and capsid_S, generated using the program ‘bediting’, was used as a reference. These maps were used as starting models for high resolution maps. The movies collected from 200 keV Glacios cryo-TEM were fed in the cryoSPARC software package and frames were aligned using Patch Motion Correction (8). The aligned frames providing one micrograph for each movie were utilized for CTF estimation using Patch CTF. A similar approach, described above, was adapted to generate high resolution maps. The 3D maps computed earlier using BSOFT were employed as initial reference maps, and the final reconstructions were carried out using the boxed particles selected after a few rounds of iterations. The density maps were visualized using ChimeraX (6). A total of 79468 phage capsids were computationally boxed out from the aligned cryo images collected on Glacios cryo-TEM. The particles were classified primarily in three groups capsid_L (59457 capsids), capsid_S (17689) and capsid_EL (2322). For capsid_L and capsid_S, further selection for better quality of particles were carried out by performing several iterations of 2D classification. This is followed by a 3D classification where reference maps were generated from the low-resolution ones (computed from the TF20 TEM datasets) with sizes computationally increased and decreased by 1% and 2% in size. These size-altered maps were used as references to further classify particle sets. The final set of particles were utilized to compute high resolution maps. The resolution of the final maps of capsid_L (3.8Å), capsid_S (4.17Å) & capsid_EL (11.93Å) at fsc cut-off at 0.143 are shown in Fig. S1-3, respectively. All the capsid internal volume determination is done in UCSF Chimera and ChimeraX (6).

### Model building and refinement

Our capsid model was created as follows. The initial atomic models for the hexon and the penton were created using Alphafold2 with the amino acid sequence of mature gp23 in which the first 120 residues were truncated (2,9). The monomers from each were fitted onto each region of the maps and symmetrized. Then, Rosetta Relax and Phenix Real-space refinement were performed for the asymmetric unit for each T=13 and T=7 along with surrounding chains (10,11). The unsupported portions of the model from the map, based on Phenix Comprehensive validation were removed.

To model the DarA and Hdf appendages in our capsid_L density, we used a template-based fitting strategy based on the P1 capsid structure from PDB 9UKM. First, the capsid portion of 9UKM was superimposed onto our reconstructed capsid model. During this alignment, the DarA and Hdf coordinates present in 9UKM were transformed simultaneously. After initial placement into the corresponding densities at the 5-fold vertices, both DarA and Hdf were further refined using the Rosetta Relax protocol within the local density environment. During this alignment, the DarA and Hdf coordinates present in 9UKM were transformed simultaneously. After initial placement into the corresponding densities at the 5-fold vertices, both DarA and Hdf were further refined using the Rosetta Relax protocol within the local density environment.

